# Cell-Type-Resolved Isoform Atlas of Human Tissues Reveals Age and Alzheimer’s Disease-Associated Splicing Changes

**DOI:** 10.1101/2025.11.14.688520

**Authors:** Ryo Yamamoto, Ting Fu, Elaine Huang, Anna Fraser, Nazim Mohammad, Xinshu Xiao, Noah Zaitlen

**Affiliations:** University of California, Los Angeles

## Abstract

Alternative splicing is a key mechanism for transcriptomic diversity, but how isoforms map to specific cell types in bulk tissues remains unclear. We present Sciege, a multimodal method that integrates bulk short-read RNA-seq with single-cell and long-read data to estimate cell type-specific isoform distributions. Through simulations, we demonstrate that Sciege accurately estimates isoform abundances and identifies differentially abundant transcripts through statistical tests. Applied to seven tissues in GTEx and brain tissue in ROSMAP datasets, Sciege generates a first-to-date multi-tissue isoform atlas and reveals isoform changes linked to cell types, aging, and Alzheimer’s disease. Validation with external cohorts and experimental data confirms our findings. Notably, we identify upregulation of the MAPT-010 isoform in AD inhibitory neurons, consistent with known methylation signatures. Our approach demonstrates the value of integrating RNA-seq data to study cell type-specific splicing and provides a foundation for further genetic and functional studies of alternative splicing across biological contexts.

## Introduction

Alternative splicing (AS) is a regulatory mechanism that generates multiple transcript isoforms from a single gene and is observed in up to 95% of multi-exon genes in the human genome^1–3^. These isoforms differ in sequence and function, contributing to diverse cellular processes^3,4^. Notably, about 15% of hereditary diseases have been linked to differential isoform usage^2,5,6^. Early investigations employing bulk RNA-seq effectively characterized AS events within well-established biological contexts, underscoring the tissue-specific nature of alternative splicing^1,7–9^. However, the lack of large scale isoform-level quantification at cell type-specific resolution is a roadblock to advance understanding of transcriptomic complexity and its relevance to disease.

Short-read RNA-seq has been widely used to characterize alternative splicing across tissues, enabling large-scale studies of splicing junction usage. However, its limited read length constrains the reconstruction of full-length transcript structures^10,11^. The introduction of long-read RNA-seq has provided a means to directly capture full-length transcripts without the need for fragmentation, substantially improving isoform-level resolution^12^ ^13–15^. Despite these advantages, most long-read RNA-seq studies to date have been performed on bulk samples, with only a few extending to single-cell assays^16–21^.

Single-cell RNA-seq (scRNA-seq) or single-nucleus RNA-seq (snRNA-seq) has yielded key insights into cell type composition, lineage relationships^22,23^, and the cellular underpinnings of complex diseases^24–27^. However, isoform-level analyses remain limited when using short-read scRNA-seq due to its pronounced 3’ or 5’ bias and limited read length^28,29^. While a few methods have been proposed to infer alternative splicing in single-cell datasets, these methods cannot fully overcome the constraints of short-read data^28,30,31^. Recently, long-read scRNA-seq approaches have been developed to directly resolve transcript isoforms in individual cells^16,17,19–21^. Widespread adoption of these protocols to population-scale has been limited by experimental cost and throughput. In contrast, large-scale efforts such as GTEx, OneKOneK and ROSMAP^32–34^ have generated extensive short-read bulk or scRNA-seq datasets, with increasing incorporation of long-read bulk data.

Here, we introduce Sciege, an integrative framework to infer cell type specific isoform distributions from broadly available short-read bulk RNA-seq data in combination with a modest number of short-read scRNA-seq and bulk long-read RNA-seq datasets from the same tissue type (Fig. 1A). Through extensive simulations and replications using external cohorts, we demonstrate that Sciege obtains robust estimation of cell type isoform abundances at population mean level. We applied Sciege to seven human tissues in the GTEx consortium and to brain tissues in the ROSMAP cohort across 21 cell types^32,33,35,36^, creating the first to date multi-tissue multi-cell type isoform abundance map. We then develop a differential testing framework and use it to examine cell type-resolved isoforms usage across aging and Alzheimer’s disease (AD), giving insights into cell type dependent regulation of splicing.

**Figure 1:**
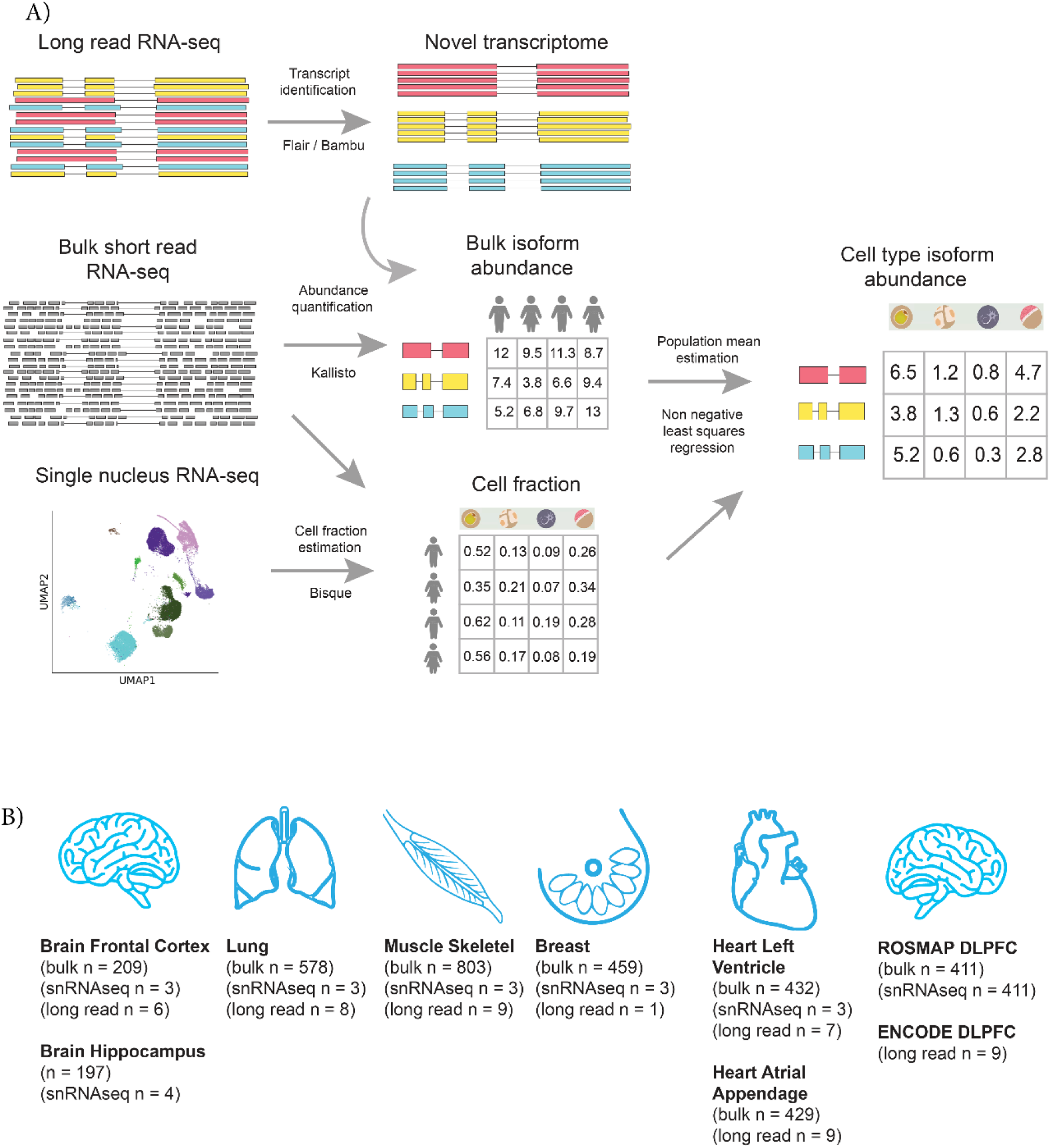
Overview of the Sciege method: A) Schematic of the Sciege workflow integrating short-read bulk RNA-seq, short-read snRNA-seq and bulk long-read RNA-seq. Subpanels depict input data types and intermediate outputs; arrows indicate analytical steps and associated methods. B) Illustration of tissues analyzed in this study. Labels indicate tissue name, data source, and sample size.

## Results

### Overview of the method

We developed Sciege, a multi-modal integration method that estimates isoform abundances at the cell type level by leveraging three data modalities, bulk short-read RNA-seq, short read scRNA-seq and bulk long-read RNA-seq, all assumed to originate from the same tissue type. We begin by identifying novel transcripts from long-read RNA-seq. Transcript discovery using bulk long-read RNA-seq is relatively well established, with several available tools^13,15,37–39^. For GTEx samples, we directly extracted transcripts already assembled by the GTEx consortium (using Flair)^15^, while for AD brain samples from ENCODE, we used Bambu^15,35,37,40,41^. These novel transcripts were combined with annotated transcripts from Gencode^42^ to construct a comprehensive reference transcriptome. We then quantified individual specific transcript abundances from bulk short-read RNA-seq samples using Kallisto, which performs pseudo-alignment of short reads to the reference transcriptome^43^. Transcript abundance quantification using bulk short-reads has been shown to yield reliable estimates for annotated transcripts, previous benchmark study based on spike-in transcripts showing more than 95% correlation across tools^44^.

We next estimate cell type composition with Bisque^45^, applying it to gene-level expression matrices derived from bulk short-read RNA-seq. We used short-read snRNA-seq data to define reference distributions of cell type specific expression. For downstream analysis, we restricted to cell types that constitute at least 3% of the total cell proportion as determined from the reference scRNA-seq data.

Finally, using the estimated cell type fractions and bulk transcript abundances, we applied non-negative least squares regression to infer then mean transcript abundance for each cell type (see Methods). We further use bootstrapping to estimate standard errors of the estimates. Sciege was applied to seven tissues from the GTEx consortium and to dorsolateral prefrontal cortex (DLPFC) samples from the ROSMAP and ENCODE datasets (Fig.1B).

### Sciege performs robust quantification of transcript abundance for major cell types across simulations

We first evaluated the performance of Sciege using realistic simulation scenarios to reflect key sources of noise in bulk RNA-seq and cell type fraction estimates (see Methods). Each simulation varied the levels of noise in bulk transcript abundance and cell type fraction estimates, two factors that directly affect the performance of cell type-specific isoform quantification. As expected, the accuracy of our method improved as the simulated levels of noise decreased, i.e., as the estimates of bulk abundance and cell type fraction correlated more with the ground truth (Fig. 2A, B). In a simple simulation setup where each gene’s expression is independent, we observed that accuracy improved with increasing sample size, stabilizing beyond approximately 50 samples (Fig. 2A). In more realistic simulations incorporating covariance between cell types and isoform abundances, accuracy was more strongly influenced by the quality of cell type fraction estimates (Fig. 2B). In both settings, the accuracy of the bulk abundance estimate was crucial in determining the accuracy of Sciege. To assess a typical level of noise in this estimate, we compared transcript abundances derived from matched short-read and long-read RNA-seq in the same GTEx individual. Focusing on novel transcripts identified from GTEx long-read data, we observed a strong positive correlation between the two platforms (Pearson R = 0.67, two-sided P < 2.2e-16, Sup Fig. 1A). This provides an empirical estimate of noise inherent in short-read quantification, supporting the use of these values as realistic parameters in our simulations (indicated in dashed line in Fig. 2A, B). In addition, it is not known whether cell type decomposition tools can be applied to whole tissue using a few numbers of single-nuclei RNA-seq samples as a reference. We tested whether Bisque performs well in this setup using leave one out cross-validation. For samples where both bulk RNA-seq and snRNA-seq are available, we computed the estimated cell proportion of one sample using bulk RNA-seq and other snRNA-seq as a reference and proceeded to compare the proportion with snRNA-seq. This validation yielded a highly concordant cell type proportion compared to ground truth (Pearson R = 0.8, Sup Fig.1B). Given this cross-validated accuracy of 0.8 for Bisque (indicated in dashed line in Fig. 2B), Sciege achieves around 70% accuracy in this simulation.

**Figure 2:**
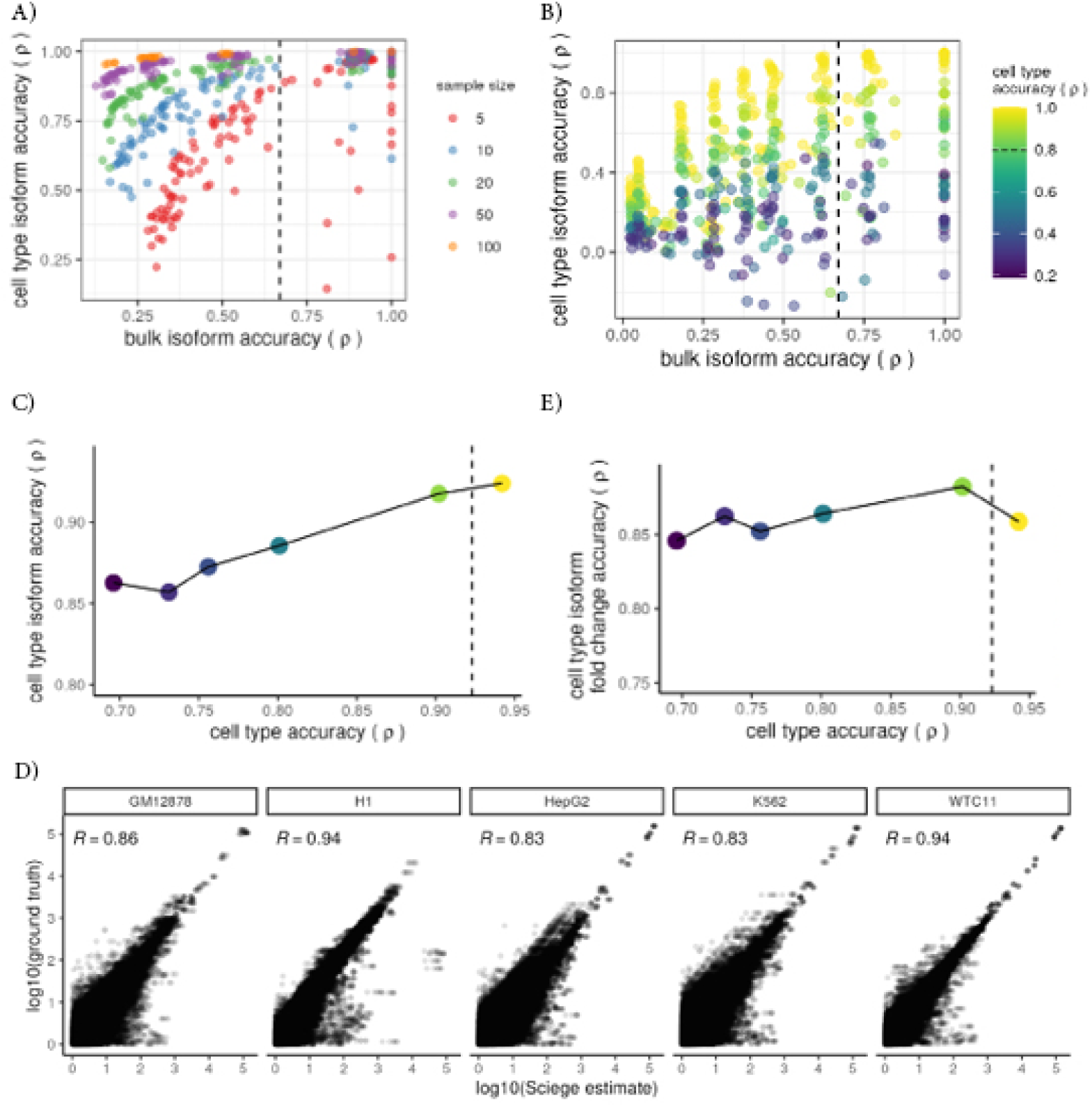
Sciege enables robust quantification of cell type-resolved transcript abundances in simulations: A) Performance of Sciege in numerical simulations with varying levels of noise in bulk transcript abundances, cell fraction estimates, and sample size. Y-axis represents the mean Pearson correlation between estimated and ground truth isoform abundances across all isoforms and cell types. X-axis shows the correlation between estimated and true bulk isoform expression. Color represents sample sizes being used in the simulation. Dashed vertical line represents x = 0.67, correlation between long-read and short-read abundance estimate. B) Same as A), but with simulation setup where isoform to isoform covariance is taken into account (See Methods). Color indicates correlation between estimated and true cell type fractions. C) Performance of Sciege in read-based simulations using mixed RNA-seq reads from five human cell lines. Y-axis shows the mean correlation between estimated and ground-truth isoform abundances across cell types; X-axis and color represent correlation between estimated and true cell type fractions. Dashed vertical line represents x = 0.923, previously reported Bisque accuracy. D) Scatter plots showing correlation between Sciege estimates and ground truth isoform abundances across all read-based simulation setups. Axes show log10-transformed expression values. Each facet corresponds to a different cell line. Pearson correlation is reported in the upper left of each panel. E) Same as C), but Y-axis shows correlation between estimated and true fold changes in isoform expression between cell types for each transcript.

We next evaluated Sciege using raw read-level data to assess its performance under realistic sequencing conditions (Sup Fig. 2A). To this end, we performed a simulation in which RNA-seq reads from five human cell lines (H1, GM12878, HepG2, K562 and WTC11) were mixed in varying proportions to simulate bulk RNA-seq samples (N = 100). Each cell line was randomly assigned to a brain cell type and we used cell type proportions derived from ROSMAP snRNA-seq to guide the mixing, mimicking realistic cellular compositions. Reads were sampled and aggregated according to these proportions to generate simulated bulk RNA-seq fastq files for 100 synthetic samples. We then applied Sciege to these read-level mixtures to estimate cell type-resolved isoform abundances. The isoform abundance in each cell line was used as ground truth, allowing direct evaluation of estimation accuracy. Sciege showed strong agreement with ground truth across cell types (Fig. 2C, D) and at varying levels of cell type fraction noise (mean R = 0.88, Fig. 2C). While cell type fraction estimates still affect the performance, their impact was reduced relative to the above numerical simulations. When accounting for realistic accuracy (indicated in dashed line, Fig. 2C), Sciege achieves more than 90% accuracy in read-based simulation. Lastly, we investigated whether relative isoform usage between pairs of cell types were accurately estimated, since these measurements would directly affect identification of cell type-distinctive transcripts. We observed that the fold changes inferred by Sciege were well correlated with true differences (Fig. 2E).

Furthermore, we validated Sciege-derived isoform abundances to those from an external single-nucleus long-read RNA-seq dataset. Using snISOr-seq data from human brain cortex tissue^20^, we observed a significant correlation between Sciege estimates and isoform abundances derived directly from the long-read snRNA-seq data (Pearson R = 0.41, Sup Fig. 2B). Notably, the correlation across samples within the snISOr-seq dataset itself was modest (Pearson R = 0.77, Sup Fig. 2C), reflecting biological and technical variability. Together, these results demonstrate that Sciege provides accurate estimates of population-level cell type-resolved isoform abundances, provided that cell fraction and bulk abundance estimates are sufficiently accurate.

### Characterization of cell type-resolved isoform abundances in human tissues

To assess isoform abundances across cell types in human tissues, we applied Sciege to seven tissues from the GTEx consortium. In total, we quantified isoform abundances for 68,810 isoforms in 21 cell types. We sought to identify isoforms that are selectively expressed in specific cell types. To do this, we developed a statistical framework that uses bootstrapped standard errors to quantify uncertainty in isoform abundance estimates and compute Z-scores for cell type-distinctive enrichment (see Methods). Specifically, we performed 100-fold bootstrapping to compute the standard error of our estimates. We then calculated Z-score comparing each cell type’s mean isoform abundance to the average across all other cell types. Transcripts were called as cell type-distinctive based on Z-score significance, fold change, and absolute expression difference. In the prior read-based simulations, this approach demonstrated high precision in identifying cell type-distinctive isoforms (Sup Fig. 3A). We also investigated whether isoform-level signals recovered known cell type marker genes. Using Cell Marker 2.0, a curated database of cell type marker genes across cell types and tissues, we found that Sciege called cell type-distinctive isoforms for 89.6% of marker genes cataloged in the database, though recovery rates varied across cell types (Sup Fig. 3B).

Applying this approach to seven tissues in the GTEx consortium, we identified a total of 46,821 cell type-distinctive isoforms across tissues. The highest number of distinctive isoforms was observed in fibroblasts and vascular endothelial cells - two cell types known for their transcriptional heterogeneity and diverse subtypes (Fig. 3A)^46,47^. We next examined whether there is a difference between cell type distinctiveness calls at the gene level and isoform level. Using the same GTEx bulk RNA-seq samples, we applied our pipeline to gene-level expression data. This analysis revealed 18,736 additional gene-cell type pairs identified as distinctive at the isoform level but not at the gene level, underscoring the added resolution of isoform-level analysis (Fig. 3B). We next assessed the consistency of cell type-resolved estimates across tissues. Estimated isoform abundances for shared cell types were highly correlated between two brain regions, supporting the robustness of Sciege’s estimates across distinct but related samples (Fig. 3C).

**Figure 3:**
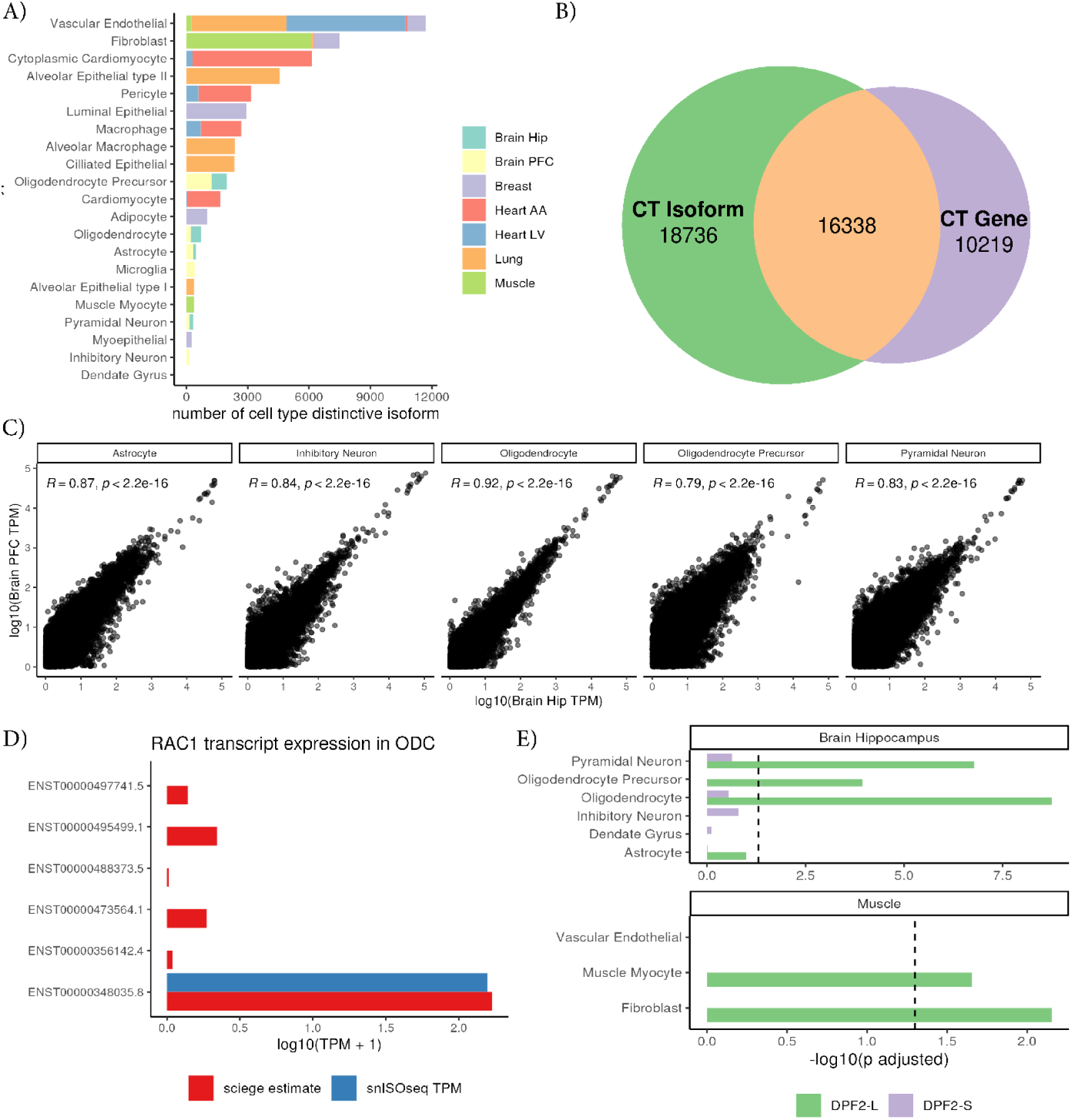
Identification of cell type-distinctive isoforms across human tissues. A) Number of cell type-distinctive isoforms identified per cell type. X-axis represents the number of distinctive isoforms. Y-axis lists cell types. Bars are colored by tissue of origin. B) Venn diagram comparing the number of genes detected as cell type-distinctive at the isoform level (left) and gene level (right) for each cell type. Overlapping regions represent gene-cell type pairs detected at both levels. C) Scatter plot showing correlation of isoform abundance estimates between hippocampus and prefrontal cortex. Axes show log10 transformed transcript TPM values. D) Isoform abundance of the *RAC1* gene in oligodendrocytes. Y-axis lists transcript IDs; X-axis represents the log-transformed TPM + 1 value of isoform expression. Blue color represents isoform abundance estimated from snISOr-seq data. Red color indicates Sciege estimated isoform abundance. E) Isoform expression of the *DPF2* gene in hippocampus and muscle tissues. X-axis shows -log10-transformed adjusted p-values. Y-axis shows cell types. Bars represent exon 7-excluded (*DPF2-S*, purple) and exon 7-included (*DPF2-L*, green) isoforms. Panels correspond to different tissues.

Moreover, we highlight a few examples in which Sciege-derived isoform abundance is validated by the long-read snRNA-seq (snISOrseq) dataset in frontal cortex (Fig. 3D, Sup Fig. 3C)^20^. For example, the *RAC1* gene was reported to have five isoforms expressed in oligodendrocytes, but only the canonical transcript ENST00000348035.6 was detected in the snISOr-seq data. Sciege correctly identified this dominant isoform, with expression patterns concordant between the two datasets (Fig. 3D). A similar result was observed for *ATP2A2*, a gene previously implicated in behavioral disorders^48^. Although other *RAC1* and *ATP2A2* isoforms had relatively similar structure to the canonical isoforms (Sup Fig. 3D), Sciege effectively recapitulated the predominant isoform usage, highlighting the effectiveness of our approach.

We also investigated whether Sciege could detect known and experimentally verified alternative splicing events. A previous study using iPSC-derived neurons showed that reduced expression of *PTBP1* promotes exon 7 inclusion in *DPF2*, a gene encoding for chromatin remodelling complex^49^, resulting in expression of the long isoform *DPF2-L* in mouse brain and muscle tissues^49^. Consistent with these findings, Sciege revealed elevated expression of *DPF2-L* in muscle and brain cell types, with limited expression of the shorter *DPF2-S* isoform (Fig. 3E).

### Cell type-resolved analysis reveals transcriptomic signatures of aging

To investigate transcriptomic changes associated with aging, we conducted differential isoform analysis using Sciege. Instead of computing population-level estimates across all samples, we stratified individuals into two age groups - young (<50 years) and old (≥50 years) - and estimated transcript abundances and corresponding standard errors within each group. For each transcript and cell type, we calculated Z-scores to compare expression between age groups. Using this approach, we discovered 2,404 age-associated transcripts across 21 cell types (Fig. 4A). Pyramidal neurons showed the highest number of differential transcripts, followed by alveolar epithelial type II cells and cytoplasmic cardiomyocytes. Despite limited sample size compared to other tissues (N = 209, Fig. 1A), prefrontal cortex tissue exhibited the highest number of age-associated transcripts among tissues analyzed (Sup Fig. 4A). Most age-associated transcripts were unique to one tissue, with minimum overlap across tissue types (Sup Fig. 4B), consistent with previous findings that transcriptomic aging signatures are specific to subregions within the same organ system^50^.

**Figure 4:**
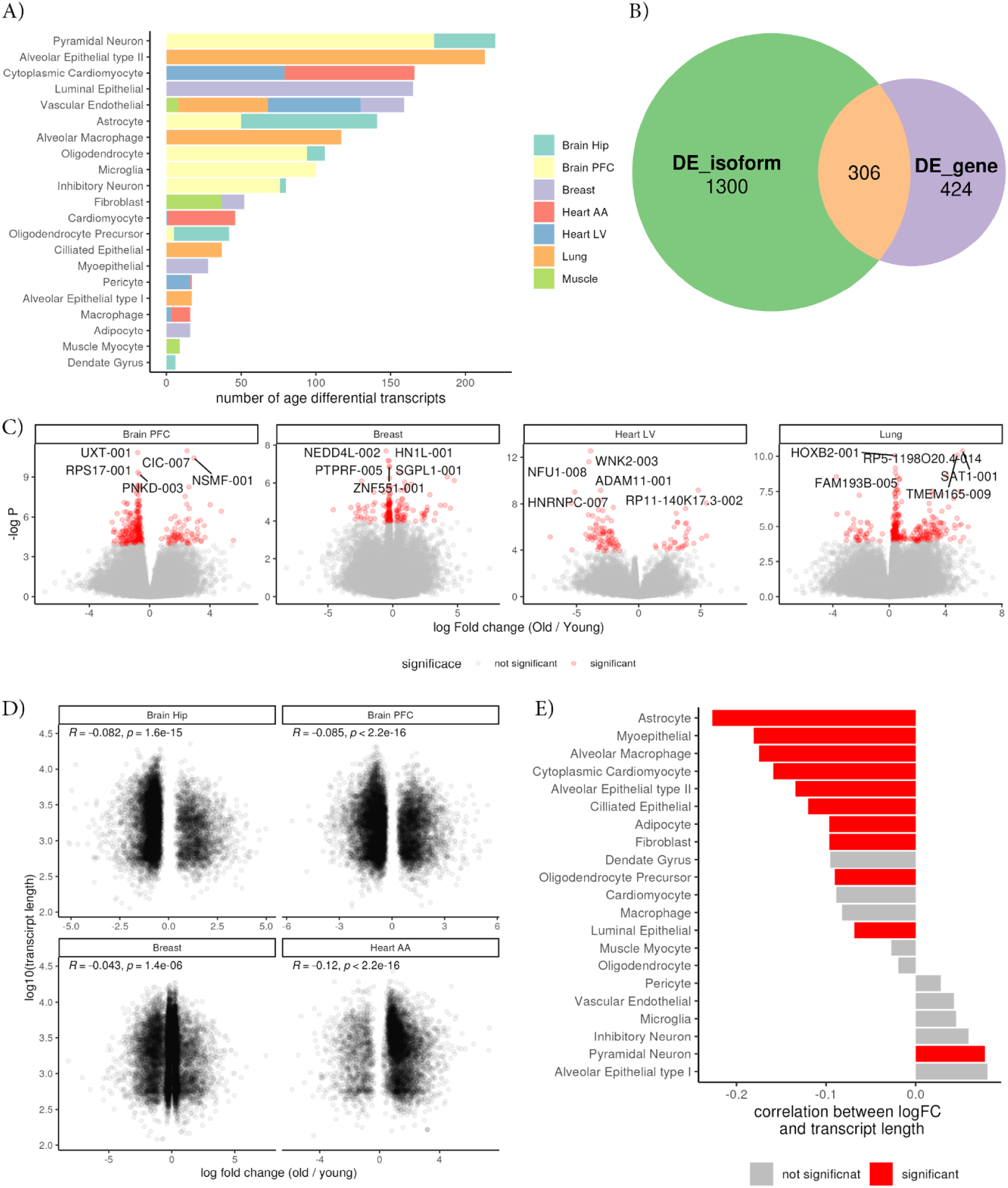
Cell type-resolved age-associated transcript changes reveal known transcriptomic features of aging: A) Number of age-associated transcripts per cell type. X-axis represents the number of differentially expressed transcripts. Y-axis lists cell types. Bars are colored by tissue of origin. B) Comparison of age-associated gene detection at the isoform level (left) vs. gene level (right). Overlapping regions indicate gene-cell type pairs identified in both analyses. C) Volcano plots showing age-associated fold change and p-value for transcripts in select tissues. X-axis represents log2 fold change (old vs. young). Y-axis represents -log10 of p-values. Color represents FDR significance. Names of a few top transcripts are shown. D) Scatter plot comparing transcript length and age-associated fold change in selected tissues. X-axis represents log2 fold change. Y-axis represents log10-transformed transcript length. Spearman correlation coefficients and p-value are shown in each panel. E) Spearman correlation between age-associated fold change and transcript length in distinct cell types. X-axis represents the correlation values. Y-axis lists cell types. Bars are colored by bonferroni corrected correlation significance (two-sided P < 2.38e-3).

We next investigated whether isoform-level analysis provides additional sensitivity beyond gene-level analysis. Across tissues, we observed a more than 3-fold increase in the number of novel age-associated genes detected at the isoform-level, underscoring the importance of isoform-resolved analysis for aging studies (Fig. 4B). Representative examples of differentially expressed isoforms are shown for selected tissues (Fig. 4C).

To assess the functional relevance of these age-associated transcripts, we performed gene set enrichment analysis using Hallmark gene sets. Enriched pathways included myogenesis, MYC targets, and immune response - gene sets previously implicated in aging (Sup Fig. 5)^51^ . Notably, we identified the *NSMF* gene in the prefrontal cortex, which has been reported to have age-associated alternative splicing in the mouse brain^52^. We also observed age-associated splicing changes in *CIC* in the prefrontal cortex, a transcriptional repressor implicated in neurological disorders^53^, consistent with earlier reports in the same GTEx tissue^54^.

Previous studies have shown that age-dependent changes in transcript expression are associated with transcript length, with longer transcripts tending to decrease in expression with age^55^. To investigate this phenomenon in our dataset, we computed the Spearman correlation between age-associated fold changes (old/young) and transcript lengths for each tissue. Consistent with earlier reports, we observed a predominant negative correlation between transcript length with age-related expression changes across tissues (Fig. 4D)^55^.

We next extended this analysis to the cell type level by calculating Spearman correlations separately for each cell type. This revealed a consistent trend of negative correlation across many cell types in human tissues, mirroring findings previously observed in mouse tissue studies (Fig. 4E)^56^. Importantly, cell type-resolved analysis enabled more granular insights. For example, astrocytes showed a strong negative correlation between transcript length and age-associated expression changes, whereas other brain cell types exhibited little or no correlation. These results suggest that transcript length-associated aging effects observed in bulk tissue may be driven by specific cell populations within those tissues.

### Cell type-resolved analysis reveals isoform-level dysregulation in Alzheimer’s disease

Aberrant alternative splicing has been implicated in AD, yet isoform-level comparisons between disease and control conditions remain limited. To study splicing alterations in AD, we applied our differential abundance framework to the ROSMAP dataset, consisting of short-read bulk and snRNA-seq data from over 400 DLFPC samples from individuals with AD and controls^33^. We identified 70 differentially expressed genes at isoform level between AD and control samples, predominantly in neuronal cell types (Sup Fig. 6A). Among these were known AD-relevant genes, such as *MAPT*, *MAPK12*, *ZHX3* and *MIR99B* (Fig. 5A)^57–60^. In addition to previously reported genes, we discovered several differentially expressed isoforms not previously implicated in AD, such as NANOS3, an RNA binding protein which has been shown to be overexpressed in cancer^61^. Z-scores for differential isoform expression of *MAPT* and *MAPK12* are shown in Fig. 5B and 5C, respectively.

**Figure 5:**
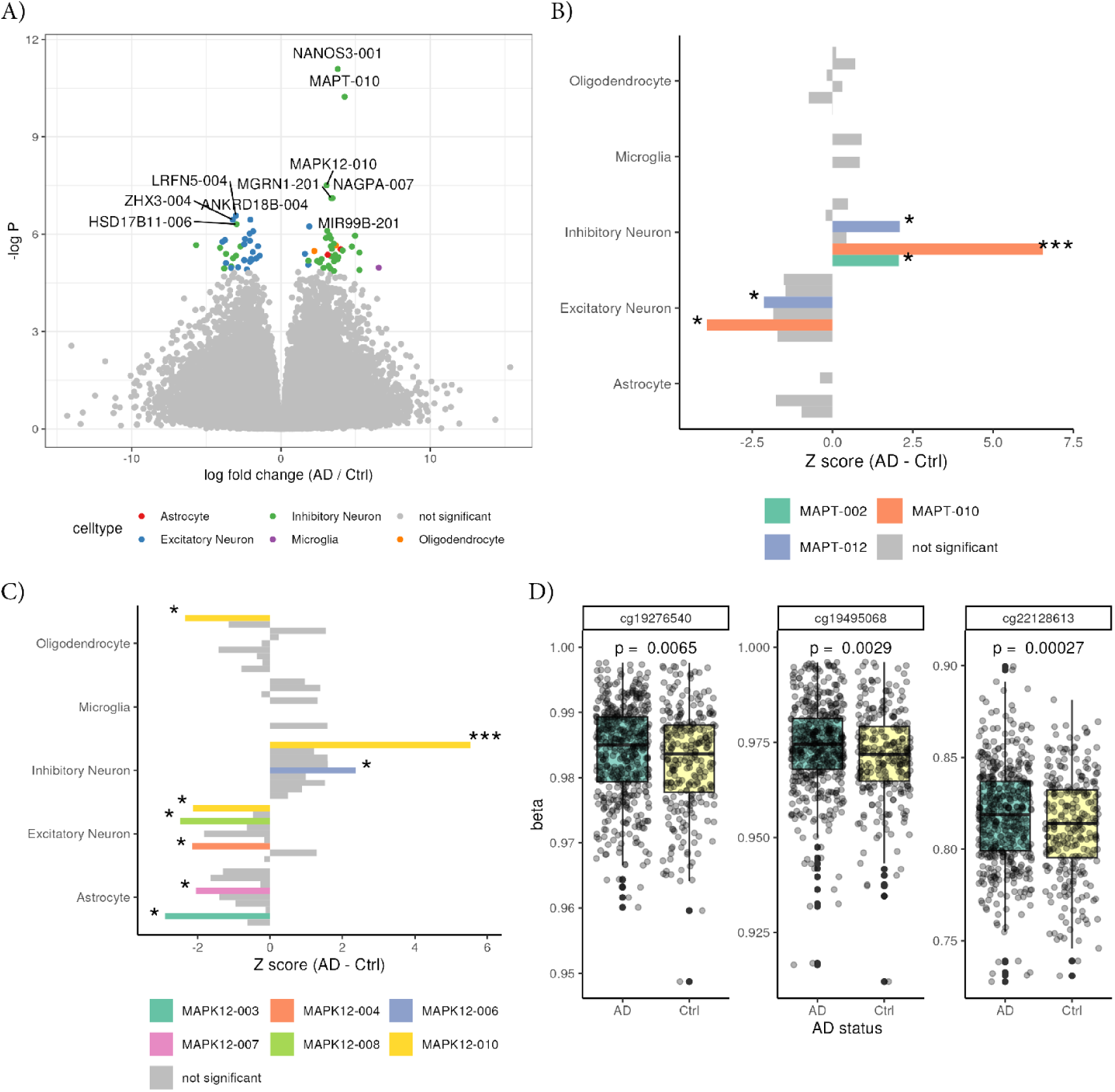
Cell type-resolved differential transcript analysis reveals isoform-specific alterations in AD. A) Volcano plot of differentially expressed transcripts between AD and control samples. X-axis shows log2 fold change (AD/control).Y-axis represents -log10 p-values. Transcripts passing FDR significance are colored by cell type; non-significant transcripts are shown in grey. Top 10 transcripts by significance are labeled. B) Z-scores for differential expression of *MAPT* isoforms across cell types. X-axis shows Z-scores comparing AD and control; Y-axis lists cell types. Bars are colored by transcript ID. Nominally significant transcripts (P < 0.05) are highlighted in color; asterisks indicate significance (*: P < 0.05, ***: FDR < 0.05). C) Same as B) but for *MAPK12* isoforms. D) Box plots showing significantly hypermethylated CpG sites in CpG21 of the *MAPT* gene. X-axis and box color indicate AD status. Y-axis shows methylation beta values. Each facet corresponds to a CpG site. P-values from linear models are shown above each plot. Centerline of the box plot indicates median, box limit indicates first and third quartiles and points indicate outliers.

Notably, we observed upregulation of the *MAPT-010* isoform in inhibitory neurons of AD brains, a small alternative transcript of the *MAPT* gene (Fig. 5B). This isoform includes one of the two CpG islands in the *MAPT* gene called CpG 21, and may contribute to the regulation of major *MAPT* isoforms^62^. Interestingly, we observed H3K27Ac peaks around 5kb upstream of the *MAPT-010* transcript, which might function as a distal promoter for this alternative transcript (Sup Fig. 7).

To further explore epigenetic regulation of *MAPT-010*, we examined cell type-specific DNA methylation patterns from a published dataset of human brain methylomes^63^. We found significant gene body hypermethylation in inhibitory neurons compared to excitatory neurons (Sup Fig. 6B). This result aligns with previous belief that hypermethylation in exonic parts helps facilitate splicing regulation^64^. We next analyzed the methylation level of CpG 21 using ROSMAP bulk methylation array data. We observed that 3 CpG sites within the island were significantly hypermethylated in AD, with 6 additional sites showing hypermethylation trends (Fig. 5D, Sup Fig. 6C). Together, these findings suggest that hypermethylation at CpG 21 may contribute to disease-associated upregulation of *MAPT-010*, potentially implicating this isoform in AD-related dysregulation.

## Discussion

Cell type-resolved analysis of alternative splicing provides important insights into how cellular contexts shape post-transcriptional regulation. In this study, we developed Sciege, a computational framework that integrates short-read bulk RNA-seq, and a small number of reference short-read scRNA-seq and bulk long-read RNA-seq to estimate transcript isoform abundances for each cell type. Through both numerical and read-based simulations, we demonstrated that Sciege provides accurate cell type-resolved isoform abundance and reliably identifies cell type-distinctive isoforms. Applying Sciege to GTEx data, we identified 46,821 cell type-distinctive isoforms, revealing widespread cell type-level splicing regulation. By estimating population-level transcript abundance per cell type, Sciege enables effective reuse of large-scale bulk short-read RNA-seq datasets while incorporating structural information from long-read data and cellular resolution from single-cell data. We further extended the framework to differential analysis, identifying isoform-level changes associated with aging and Alzheimer’s disease in a cell type–resolved manner, including known and novel transcriptomic features.

Despite its utility, our approach is built on several assumptions that warrant consideration. First, Sciege assumes that cell type fraction estimates from deconvolution are accurate. Our numerical simulations show that errors in cell fraction estimates have a greater impact on isoform quantification than errors in bulk abundance estimates. While we observed good agreement between estimated and expected cell fractions in our datasets, the choice of deconvolution tool and availability of appropriate single-cell reference panels will influence performance. In this study, we used Bisque, which assumes that matched bulk and single-cell samples are available^45^. In future applications, alternative tools may be preferable when using external reference panels or unmatched data.

It is well established that correcting for covariates improves the accuracy and interpretability of differential expression analyses. In our current framework, we assume that potential confounders affect stratified groups (e.g., age or disease status) in similar ways. However, this assumption may not hold in cases involving complex interactions, such as the interplay between age and AD status, which could introduce bias into the results. While our method enables robust estimation of population mean isoform abundance per cell type, it does not currently allow for individual-level modeling or covariate adjustment. Future studies that incorporate individual-level deconvolution of isoform abundances will be necessary to address this question.

Genetic regulation of alternative splicing plays a critical role in human diseases, with splicing events in some cases exerting stronger phenotypic effects than gene expression^65,66^. Cell type-specific analyses have further elucidated functions of previously unexplained genetic variants^67^. The isoform abundance estimates generated by our method could be leveraged for genetic association studies by stratifying population groups based on genotypes. However, applying this approach across the genome would be computationally intensive and limited by the need to estimate group-level differences. Again, this problem can be addressed via deconvolution of isoform abundances at the individual level. While methods for individual-level deconvolution of gene expression have been developed^68,69^, analogous tools for transcript-level resolution are currently lacking. Future studies to extend these methods to isoform-level quantification would facilitate a deeper understanding of the genetic architecture of alternative splicing at the cellular level.

In summary, Sciege enables isoform-level quantification at the level of major cell types by integrating complementary sequencing modalities. Using this approach, we generated a transcript-level atlas of isoform abundance across human tissues and identified cell type-specific transcriptomic alterations in aging and Alzheimer’s disease. These results underscore the importance of isoform-level resolution in understanding gene regulation and offer a path forward for splicing-focused studies in complex tissues.

## Methods

### Datasets

Raw short-read and long-read RNA-seq data for seven human tissues were obtained from the GTEx project (dbGaP accession: phs000424.v8.p2). Processed single-nucleus RNA-seq (snRNA-seq) data for five non-brain tissues were downloaded in h5ad format from the GTEx portal^36^. For brain tissues, processed snRNA-seq data in txt file format were obtained from the Broad Institute’s Single Cell Portal (https://singlecell.broadinstitute.org/single_cell). For the ROSMAP cohort, raw short-read bulk RNA-seq, snRNA-seq, and processed DNA methylation array data were downloaded from the AD Knowledge Portal (https://adknowledgeportal.synapse.org/). Long-read RNA-seq data from dorsolateral prefrontal cortex samples were acquired from the ENCODE project (https://www.encodeproject.org/). CpG-methylation data for excitatory neurons and inhibitory neurons were downloaded from the following link (https://cellxgene.cziscience.com/collections/fdebfda9-bb9a-4b4b-97e5-651097ea07b0).

### Identification of novel transcripts from bulk long-read RNA-seq

To identify transcripts present in the cellular context of interest, we established tools for long-read RNA-seq analysis, including Flair and Bambu^15,37^. For GTEx data, we leveraged previously processed long-read transcript annotations available from the GTEx portal, which were generated using Flair. To restrict the analysis to transcripts expressed in the tissues of interest, we applied the “Flair quantify” function to compute transcript-level coverage in each sample, and retained transcripts supported by ≥5 reads. The resulting set of filtered transcripts was used as the reference for isoform quantification in the corresponding GTEx short-read RNA-seq samples. For the analysis of Alzheimer’s disease brain tissue in the ROSMAP dataset, we used brain (AD and controls) long-read RNA-seq data from ENCODE and applied Bambu for transcript discovery^37^. Briefly, Bambu takes aligned long-read bam files as input, performs splicing junction correction and clusters reads to reconstruct transcript models. A recent benchmarking study showed that Bambu is highly accurate and robust across datasets^70^. Transcripts identified from the ENCODE long-read data were used as the reference set for transcript quantification in the ROSMAP short-read RNA-seq samples.

### Quantification of bulk transcript abundance from short-read RNA-seq

To quantify isoform expression levels in bulk short-read RNA-seq samples, we used kallisto (v0.46.1) for pseudo-alignment and transcript quantification^43^. A unified transcript reference was first constructed by combining annotated transcripts from GENCODE v26 with novel transcripts identified from long-read RNA-seq (see above). This combined transcriptome was indexed using the “kallisto index” function. For each sample, transcript abundance was quantified using the “kallisto quant” function, generating transcript per million bases (TPM) values. The resulting TPM vectors were aggregated across samples to generate a sample-by-transcript matrix for each tissue, serving as input for downstream isoform-level deconvolution and differential analysis.

### Cell type proportion estimation from bulk RNA-seq and snRNA-seq

To compute cell type proportion from bulk short-read RNA-seq for each sample, we applied Bisque in reference based decomposition mode using tissue type-matched snRNA-seq as the reference^45^. Bisque identifies cell type-specific marker genes from the raw snRNA-seq count matrix and learns a transformation between bulk gene expression and pseudo bulk single-cell profiles^45^. For GTEx tissues, we used matched bulk RNA-seq and snRNA-seq samples when available, excluding these matched samples from downstream isoform-level analysis to avoid circularity. For the ROSMAP cohort, where snRNA-seq dta were available for all individuals, cell type fractions were calculated directly from the snRNA-seq profiles based on previously published cell type annotations^71^.

### Estimation of population-mean isoform abundance per cell type

To quantify isoform abundance at the cell type level, we computed the population mean expression of each transcript by integrating bulk transcript abundance and estimated cell type proportions. For each transcript, we employed non-negative least square (NNLS) regression model of the form:

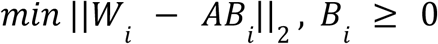

where *W_i_* is a vector of bulk expression values of transcript *i* across samples, *A* denotes the matrix of estimated cell type fractions, *B_i_* represents the population mean abundance of transcript *i* for each cell type. This formulation assumes that the observed bulk expression for each transcript can be modeled as a linear combination of cell type–specific expression, weighted by the estimated cell type fractions. We used the nnls package in R to solve the NNLS optimization for each transcript, along with the boot package to perform 100 rounds of bootstrapping and compute standard errors for each estimate^72–74^.

### Numerical simulation setup

To assess the robustness and accuracy of our method, we designed a series of simulation studies comprising three distinct settings, namely pure numerical simulation, real parameter-based numerical simulation and read-based simulation. In the pure numerical simulation, we sampled cell type fraction, bulk isoform expression as follows:

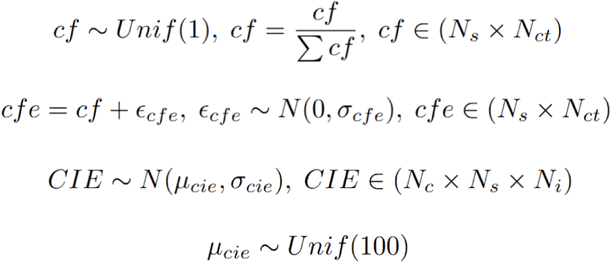

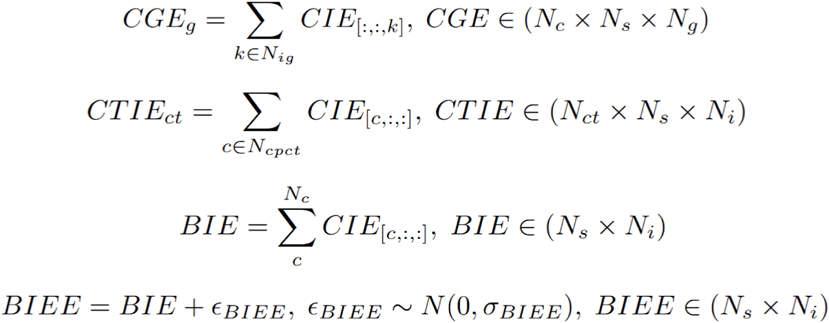

where *N_s_*denotes the number of samples, *N_ct_* denotes the number of cell types, *N_i_* denotes the number of isoforms, *N_g_*denotes the number of genes, *N_c_* denotes the number of cells, *cf* denotes the ground truth cell type fraction matrix, *cfe* denotes the estimated cell type fraction matrix (with added noise), *CIE* denotes the cell-level isoform expression 3D matrix, *CGE* denotes the cell-level gene expression 3D matrix, *CTIE* denotes the cell type-level isoform expression 3D matrix, *BIE* denotes the bulk isoform expression matrix, *BIEE* denotes the estimated bulk isoform expression matrix (with added noise).

The *BIEE* and *cfe* matrices were used as inputs to the non-negative least squares regression model to estimate cell type-level isoform expression. These estimates were then compared to the ground truth *CTIE* matrix to evaluate the performance. In this setting, isoform expression and cell type fractions were sampled independently across samples and transcripts, therefore no sample-level or transcript-level co-variation was introduced.

In the real parameter-based simulation, we introduce more realistic variation. Cell fractions were sampled from observed fractions in ROSMAP snRNA-seq data and isoform expression values (µ_cie_) were sampled from real GTEx long-read isoform data from heart left ventricle tissue. This allowed us to model biologically relevant co-variation in cell type fractions and isoform expression while still keeping noise terms independent. These simulations provide a more representative benchmark for evaluating the performance of our method under realistic parameter constraints. Across all simulations, we systematically explored combinations of parameters such as sample size, noise levels, and correlation between estimated and true bulk expression and cell fractions. All parameters and ranges used in these simulations are provided in Sup Table S1.

### Read-based simulation setup

To further approximate real-world conditions, we devised a read-based simulation framework (Fig. Sup 2). This approach aims to mimic the data characteristics encountered in typical bulk RNA-seq analyses while providing a realistic benchmark for isoform quantification. As in the above simulations, we first sampled cell type proportions from annotated snRNA-seq data in the ROSMAP cohort. Each cell type was then assigned to a human cell line, which served as the ground truth reference for isoform expression in the corresponding “cell type”. Five human cell lines were used (H1, GM12878, HepG2, K562 and WTC11) whose RNA-seq data were obtained from the ENCODE data portal. The mapping between cell types and cell lines is provided in Sup Table S2. We opted to use bulk short-read RNA-seq data from these cell lines rather than snRNA-seq data to avoid 3’ or 5’ bias in the read distribution and better recapitulate the read distribution found in conventional bulk RNA-seq.

For each simulated sample, reads were sampled from the FASTQ files of the assigned cell lines according to the sampled cell type proportions. These reads were then aggregated to construct synthetic bulk RNA-seq FASTQ files. The resulting files were quantified using kallisto to generate a bulk isoform expression matrix.

To establish the ground truth, we also quantified isoform abundances separately for each original cell line using kallisto, yielding the reference cell line–specific isoform expression profiles. The simulated bulk isoform matrix and cell type fraction estimates (with added noise) were then processed using the NNLS model to generate estimated cell type-resolved isoform abundances, which were subsequently compared to the ground truth.

### Statistical testing using population mean estimates with bootstrapping

To assess the statistical significance of cell type-resolved isoform abundances, we performed hypothesis testing using bootstrapped standard errors derived from our population mean estimates. For each transcript, we first tested whether its expression was significantly higher in one cell type compared to others. Specifically, we performed a two-sided Z-test comparing the abundance in the focal cell type against the average across all other cell types. We applied false discovery rate (FDR) correction using the Benjamini-Hochberg procedure and retained transcripts with adjusted p < 0.05. To further ensure biological relevance, we applied additional filters requiring a fold change >2 and an absolute expression difference >2 TPM, which helped eliminate transcripts with low expression but spuriously significant p-values.

For differential testing across biological conditions, we stratified samples by covariates such as age (GTEx) and Alzheimer’s disease status (ROSMAP), and estimated population mean isoform abundances separately for each group using the same NNLS framework. In the GTEx analysis, we classified samples into “young” and “old” groups using a 50-year age cutoff. In the ROSMAP dataset, AD status was determined based on CERAD scores: individuals with scores of 1 or 2 were assigned to the AD group, while scores of 3 or 4 were assigned to the control group. Sample sizes for each group are listed in Sup Table S3. We computed Z-scores for each transcript and cell type by comparing population means between the two stratified groups and applied the same FDR correction and expression-level filtering criteria as described above for the cell type–distinctiveness analysis.

### Gene set enrichment analysis with age-differential transcripts

To investigate whether age-differential transcripts we identified are enriched in age-related functions, we conducted gene set enrichment analysis using hallmark gene sets^75^. Over representation analysis for genes with age-differential transcript for each tissue are conducted using the clusterprofiler package in R^76^.

### Differential methylation testing in ROSMAP

To investigate whether CpG sites within the CpG 21 island of the *MAPT* gene are differentially methylated in AD, we analyzed bulk DNA methylation array data from the ROSMAP cohort^77^.

Differential methylation analysis was conducted using a linear regression model, following procedures described in a previous study on the same dataset^77^. The model included AD status as the primary variable of interest and adjusted for relevant covariates: study origin (Religious Orders Study [ROS] or Memory and Aging Project [MAP]), sex, postmortem interval, APOE genotype, and age of death. We tested whether AD status was significantly associated with methylation beta values at individual CpG sites, and reported effect sizes and p-values accordingly.

## Data Availability

GTEx version 8 and 9 processed data was downloaded from https://www.gtexportal.org/home/downloads/adult-gtex/bulk_tissue_expression. GTEx protected data was downloaded under dbGapaccession: phs000424.v8.p2. For brain snRNA-seq data was available in Broad Institute’s Single Cell Portal (https://singlecell.broadinstitute.org/single_cell). For the ROSMAP cohort, raw short-read bulk RNA-seq, snRNA-seq, and processed DNA methylation array data were downloaded from the AD Knowledge Portal (https://adknowledgeportal.synapse.org/). Long-read RNA-seq data from dorsolateral prefrontal cortex samples were acquired from the ENCODE project (https://www.encodeproject.org/) with following accession codes (ENCSR094NFM, ENCSR169YNI, ENCSR205QMF, ENCSR257YUB, ENCSR316ZTD, ENCSR462COR, ENCSR463IDK, ENCSR690QHM, ENCSR697ASE). CpG-methylation data for excitatory neurons and inhibitory neurons were downloaded from the following link (https://cellxgene.cziscience.com/collections/fdebfda9-bb9a-4b4b-97e5-651097ea07b0).

## Code Availability

Software implementation for Sciege is available in https://github.com/ryo1024/Sciege.

## Notes

https://github.com/ryo1024/Sciege

### Competing Interest Statement

The authors have declared no competing interest.

## References

1. Pan, Q., Shai, O., Lee, L. J., Frey, B. J. & Blencowe, B. J. Deep surveying of alternative splicing complexity in the human transcriptome by high-throughput sequencing. Nat Genet 40, 1413–1415 (2008).

2. Jiang, W. & Chen, L. Alternative splicing: Human disease and quantitative analysis from high-throughput sequencing. Computational and Structural Biotechnology Journal 19, 183–195 (2021).

3. Leoni, G., Le Pera, L., Ferrè, F., Raimondo, D. & Tramontano, A. Coding potential of the products of alternative splicing in human. Genome Biology 12, R9 (2011).

4. Wang, Y., et al. Mechanism of alternative splicing and its regulation (Review). Biomedical Reports 3, 152–158 (2015).

5. Cui, Y., Cai, M. & Stanley, H. E. Comparative Analysis and Classification of Cassette Exons and Constitutive Exons. BioMed Research International 2017, 7323508 (2017).

6. Marquez, Y., Brown, J. W. S., Simpson, C., Barta, A. & Kalyna, M. Transcriptome survey reveals increased complexity of the alternative splicing landscape in Arabidopsis. Genome Res. 22, 1184–1195 (2012).

7. Wang, E. T., et al. Alternative isoform regulation in human tissue transcriptomes. Nature 456, 470–476 (2008).

8. Raj, B. & Blencowe, B. J. Alternative Splicing in the Mammalian Nervous System: Recent Insights into Mechanisms and Functional Roles. Neuron 87, 14–27 (2015).

9. Martinez, N. M. & Lynch, K. W. Control of alternative splicing in immune responses: many regulators, many predictions, much still to learn. Immunological Reviews 253, 216–236 (2013).

10. Tilgner, H., et al. Comprehensive transcriptome analysis using synthetic long-read sequencing reveals molecular co-association of distant splicing events. Nat Biotechnol 33, 736–742 (2015).

11. Tilgner, H., Grubert, F., Sharon, D. & Snyder, M. P. Defining a personal, allele-specific, and single-molecule long-read transcriptome. Proceedings of the National Academy of Sciences 111, 9869–9874 (2014).

12. Amarasinghe, S. L., et al. Opportunities and challenges in long-read sequencing data analysis. Genome Biology 21, 30 (2020).

13. Wyman, D., et al. A technology-agnostic long-read analysis pipeline for transcriptome discovery and quantification. 672931 Preprint at 10.1101/672931 (2020).

14. Byrne, A., et al. Nanopore long-read RNAseq reveals widespread transcriptional variation among the surface receptors of individual B cells. Nat Commun 8, 16027 (2017).

15. Tang, A. D., et al. Full-length transcript characterization of SF3B1 mutation in chronic lymphocytic leukemia reveals downregulation of retained introns. Nat Commun 11, 1438 (2020).

16. Picelli, S., et al. Full-length RNA-seq from single cells using Smart-seq2. Nat Protoc 9, 171–181 (2014).

17. Ramsköld, D., et al. Full-length mRNA-Seq from single-cell levels of RNA and individual circulating tumor cells. Nat Biotechnol 30, 777–782 (2012).

18. Philpott, M., et al. Nanopore sequencing of single-cell transcriptomes with scCOLOR-seq. Nat Biotechnol 39, 1517–1520 (2021).

19. Shiau, C.-K., et al. High throughput single cell long-read sequencing analyses of same-cell genotypes and phenotypes in human tumors. Nat Commun 14, 4124 (2023).

20. Hardwick, S. A., et al. Single-nuclei isoform RNA sequencing unlocks barcoded exon connectivity in frozen brain tissue. Nat Biotechnol 40, 1082–1092 (2022).

21. Joglekar, A., et al. Single-cell long-read sequencing-based mapping reveals specialized splicing patterns in developing and adult mouse and human brain. Nat Neurosci 27, 1051–1063 (2024).

22. Treutlein, B., et al. Reconstructing lineage hierarchies of the distal lung epithelium using single-cell RNA-seq. Nature 509, 371–375 (2014).

23. DeLaughter, D. M., et al. Single-Cell Resolution of Temporal Gene Expression during Heart Development. Developmental Cell 39, 480–490 (2016).

24. Llorens-Bobadilla, E., et al. Single-Cell Transcriptomics Reveals a Population of Dormant Neural Stem Cells that Become Activated upon Brain Injury. Cell Stem Cell 17, 329–340 (2015).

25. Buettner, F., et al. Computational analysis of cell-to-cell heterogeneity in single-cell RNA-sequencing data reveals hidden subpopulations of cells. Nat Biotechnol 33, 155–160 (2015).

26. Chen, R., Wu, X., Jiang, L. & Zhang, Y. Single-Cell RNA-Seq Reveals Hypothalamic Cell Diversity. Cell Reports 18, 3227–3241 (2017).

27. Cornish, A. J., Filippis, I., David, A. & Sternberg, M. J. E. Exploring the cellular basis of human disease through a large-scale mapping of deleterious genes to cell types. Genome Med 7, 95 (2015).

28. Pan, L., Dinh, H. Q., Pawitan, Y. & Vu, T. N. Isoform-level quantification for single-cell RNA sequencing. Bioinformatics 38, 1287–1294 (2022).

29. Westoby, J., Artemov, P., Hemberg, M. & Ferguson-Smith, A. Obstacles to detecting isoforms using full-length scRNA-seq data. Genome Biol 21, 74 (2020).

30. Xiang, X., He, Y., Zhang, Z. & Yang, X. Interrogations of single-cell RNA splicing landscapes with SCASL define new cell identities with physiological relevance. Nat Commun 15, 2164 (2024).

31. Arzalluz-Luque, A., Salguero, P., Tarazona, S. & Conesa, A. acorde unravels functionally interpretable networks of isoform co-usage from single cell data. Nat Commun 13, 1828 (2022).

32. Aguet, F., et al. Genetic effects on gene expression across human tissues. Nature 550, 204–213 (2017).

33. Bennett, D. A. et al. Religious Orders Study and Rush Memory and Aging Project. Journal of Alzheimer’s Disease 64, S161–S189 (2018).

34. Yazar, S., et al. Single-cell eQTL mapping identifies cell type–specific genetic control of autoimmune disease. Science 376, eabf3041 (2022).

35. Glinos, D. A., et al. Transcriptome variation in human tissues revealed by long-read sequencing. Nature 608, 353–359 (2022).

36. Eraslan, G., et al. Single-nucleus cross-tissue molecular reference maps toward understanding disease gene function. Science 376, eabl4290 (2022).

37. Chen, Y., et al. Context-aware transcript quantification from long-read RNA-seq data with Bambu. Nat Methods 20, 1187–1195 (2023).

38. Prjibelski, A. D., et al. Accurate isoform discovery with IsoQuant using long reads. Nat Biotechnol 41, 915–918 (2023).

39. Kovaka, S., et al. Transcriptome assembly from long-read RNA-seq alignments with StringTie2. Genome Biology 20, 278 (2019).

40. Dunham, I., et al. An integrated encyclopedia of DNA elements in the human genome. Nature 489, 57–74 (2012).

41. Luo, Y., et al. New developments on the Encyclopedia of DNA Elements (ENCODE) data portal. Nucleic Acids Res 48, D882–D889 (2020).

42. Frankish, A., et al. GENCODE: reference annotation for the human and mouse genomes in 2023. Nucleic Acids Research 51, D942–D949 (2023).

43. Bray, N. L., Pimentel, H., Melsted, P. & Pachter, L. Near-optimal probabilistic RNA-seq quantification. Nat Biotechnol 34, 525–527 (2016).

44. Zhang, C., Zhang, B., Lin, L.-L. & Zhao, S. Evaluation and comparison of computational tools for RNA-seq isoform quantification. BMC Genomics 18, 583 (2017).

45. Jew, B., et al. Accurate estimation of cell composition in bulk expression through robust integration of single-cell information. Nat Commun 11, 1971 (2020).

46. Trimm, E. & Red-Horse, K. Vascular endothelial cell development and diversity. Nat Rev Cardiol 20, 197–210 (2023).

47. Page, M. L., et al. Surveying the landscape of RNA isoform diversity and expression across 9 GTEx tissues using long-read sequencing data. bioRxiv 2024.02.13.579945 (2024) doi:10.1101/2024.02.13.579945.

48. Nakajima, K., et al. Brain-specific heterozygous loss-of-function of ATP2A2, endoplasmic reticulum Ca2+ pump responsible for Darier’s disease, causes behavioral abnormalities and a hyper-dopaminergic state. Human Molecular Genetics 30, 1762–1772 (2021).

49. Nazim, M., et al. Alternative splicing of a chromatin modifier alters the transcriptional regulatory programs of stem cell maintenance and neuronal differentiation. Cell Stem Cell 31, 754–771.e6 (2024).

50. Yamamoto, R., et al. Tissue-specific impacts of aging and genetics on gene expression patterns in humans. Nat Commun 13, 5803 (2022).

51. Liberzon, A., et al. The Molecular Signatures Database (MSigDB) hallmark gene set collection. Cell Syst 1, 417–425 (2015).

52. Winsky-Sommerer, R., King, H. A., Iadevaia, V., Möller-Levet, C. & Gerber, A. P. A post-transcriptional regulatory landscape of aging in the female mouse hippocampus. Front. Aging Neurosci. 15, (2023).

53. Ruiz, I., Wiltrout, K., Stredny, C. & Mahida, S. CIC-Related Neurodevelopmental Disorder: A Review of the Literature and an Expansion of Genotype and Phenotype. Genes (Basel) 15, 1425 (2024).

54. Wang, K., et al. Comprehensive map of age-associated splicing changes across human tissues and their contributions to age-associated diseases. Sci Rep 8, 10929 (2018).

55. Stoeger, T., et al. Aging is associated with a systemic length-associated transcriptome imbalance. Nat Aging 2, 1191–1206 (2022).

56. Kimmel, J. C., et al. Murine single-cell RNA-seq reveals cell-identity- and tissue-specific trajectories of aging. Genome Res. 29, 2088–2103 (2019).

57. Strang, K. H., Golde, T. E. & Giasson, B. I. *MAPT* mutations, tauopathy, and mechanisms of neurodegeneration. Laboratory Investigation 99, 912–928 (2019).

58. Apostolakou, A. E., Sula, X. K., Nastou, K. C., Nasi, G. I. & Iconomidou, V. A. Exploring the conservation of Alzheimer-related pathways between H. sapiens and C. elegans: a network alignment approach. Sci Rep 11, 4572 (2021).

59. Shippy, D. C. & Ulland, T. K. Exploring the zinc-related transcriptional landscape in Alzheimer’s disease. IBRO Neuroscience Reports 13, 31–37 (2022).

60. Ye, X., et al. MicroRNAs 99b-5p/100-5p Regulated by Endoplasmic Reticulum Stress are Involved in Abeta-Induced Pathologies. Front. Aging Neurosci. 7, (2015).

61. Zhang, F., Liu, R., Liu, C., Zhang, H. & Lu, Y. Nanos3, a cancer-germline gene, promotes cell proliferation, migration, chemoresistance, and invasion of human glioblastoma. Cancer Cell International 20, 197 (2020).

62. Caillet-Boudin, M.-L., Buée, L., Sergeant, N. & Lefebvre, B. Regulation of human MAPT gene expression. Molecular Neurodegeneration 10, 28 (2015).

63. Tian, W., et al. Single-cell DNA methylation and 3D genome architecture in the human brain. Science 382, eadf5357 (2023).

64. Maor, G. L., Yearim, A. & Ast, G. The alternative role of DNA methylation in splicing regulation. Trends in Genetics 31, 274–280 (2015).

65. Bhattacharya, A., et al. Isoform-level transcriptome-wide association uncovers extensive novel genetic risk mechanisms for neuropsychiatric disorders in the human brain. 2022.08.23.22279134 Preprint at 10.1101/2022.08.23.22279134 (2023).

66. Hervoso, J. L., et al. Splicing-specific transcriptome-wide association uncovers genetic mechanisms for schizophrenia. The American Journal of Human Genetics 111, 1573–1587 (2024).

67. Boltz, T., et al. Cell-type deconvolution of bulk-blood RNA-seq reveals biological insights into neuropsychiatric disorders. The American Journal of Human Genetics 111, 323–337 (2024).

68. Wang, J., Roeder, K. & Devlin, B. Bayesian estimation of cell type–specific gene expression with prior derived from single-cell data. Genome Res. 31, 1807–1818 (2021).

69. Rahmani, E., et al. Cell-type-specific resolution epigenetics without the need for cell sorting or single-cell biology. Nat Commun 10, 3417 (2019).

70. Pardo-Palacios, F. J., et al. Systematic assessment of long-read RNA-seq methods for transcript identification and quantification. Nat Methods 21, 1349–1363 (2024).

71. Fujita, M., et al. Cell subtype-specific effects of genetic variation in the Alzheimer’s disease brain. Nat Genet 56, 605–614 (2024).

72. nnls: The Lawson-Hanson Algorithm for Non-Negative Least Squares (NNLS) version 1.6 from CRAN. https://rdrr.io/cran/nnls/.

73. Davison, A. C. & Hinkley, D. V. Bootstrap Methods and Their Application. (Cambridge University Press, Cambridge, 1997). doi:10.1017/CBO9780511802843.

74. boot: Bootstrap Functions (Originally by Angelo Canty for S) version 1.3-31 from CRAN. https://rdrr.io/cran/boot/.

75. Liberzon, A., et al. The Molecular Signatures Database (MSigDB) hallmark gene set collection. Cell Syst 1, 417–425 (2015).

76. Wu, T., et al. clusterProfiler 4.0: A universal enrichment tool for interpreting omics data. The Innovation 2, 100141 (2021).

77. De Jager, P. L., et al. Alzheimer’s disease: early alterations in brain DNA methylation at ANK1, BIN1, RHBDF2 and other loci. Nat Neurosci 17, 1156–1163 (2014).

