## Supplement for "Cell-Type-Resolved Isoform Atlas of Human Tissues Reveals Age and Alzheimer’s Disease-Associated Splicing Changes"

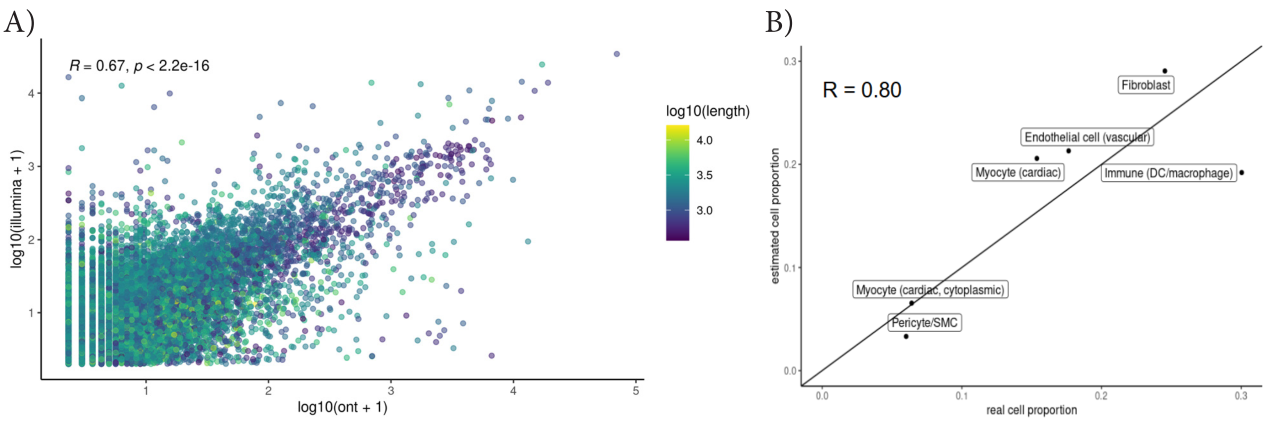


Figure S1: A) Scatter plot comparing transcript quantification between illumina short-reads and Oxford Nanopore long-reads in matching samples. Transcript used in the analysis are novel transcripts identified from same long-read previously in GTEx v9 study. X-axis represents log transformed count + 1 from long-reads. Y-axis represents log transformed count + 1 from illumina short-reads. Color scale refers to log transformed transcript length. Pearson correlation and p-values are shown on top left. B) Scatter plot comapring estimated cell type decomposition from Bisque and real cell proportion from snRNA-seq of the same sample. X-axis represents real cell proportion values from snRNA-seq. Y-axis represents estimatesd cell proportion from Bisque. Pearson correlation are shown on top left.

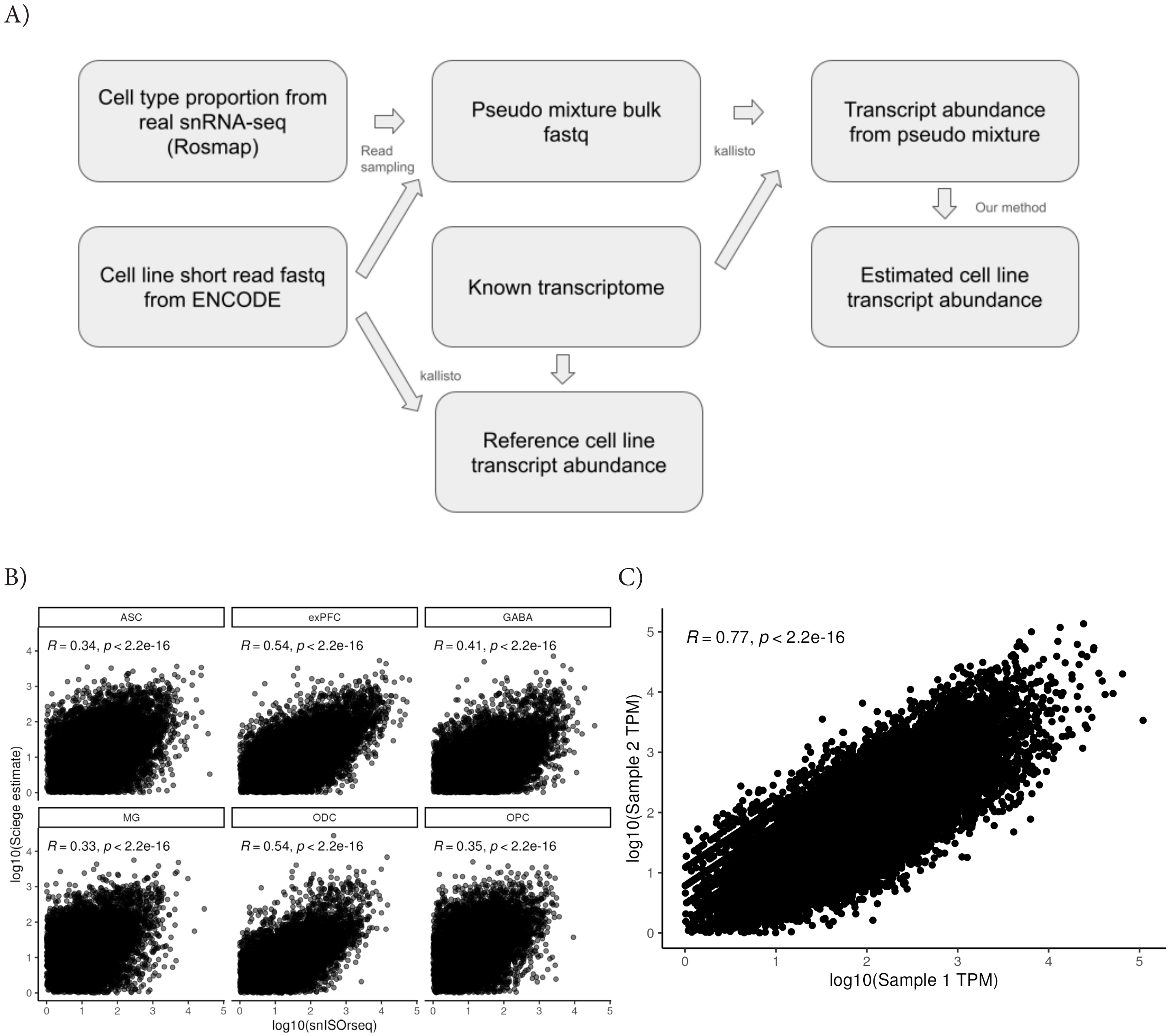


Figure S2: A) Diagram for read-level simulation. Boxes represent data and processed data. Arrows represent combinations of data and their methodology. B) Scatter plot showing correlation between our estimated abundance in GTEx prefrontal cortex tissue and single-nucleus long-read RNA-seq in brain cortex tissue. X-axis shows log-transformed TPM abudnace from snISOseq. Y-axis shows log-transformed TPM from our estimates. Pearson correlation and p-values are shown at the top. Facet shows cell type names. C) Scatter plot showing correlation between sample 1 and sample 2 from snISOrseq. X-axis and Y-axis show long transformed TPM abudnace from sample 1 and sample 2 respectively. Pearson coorelation and p-values are shown at the top.

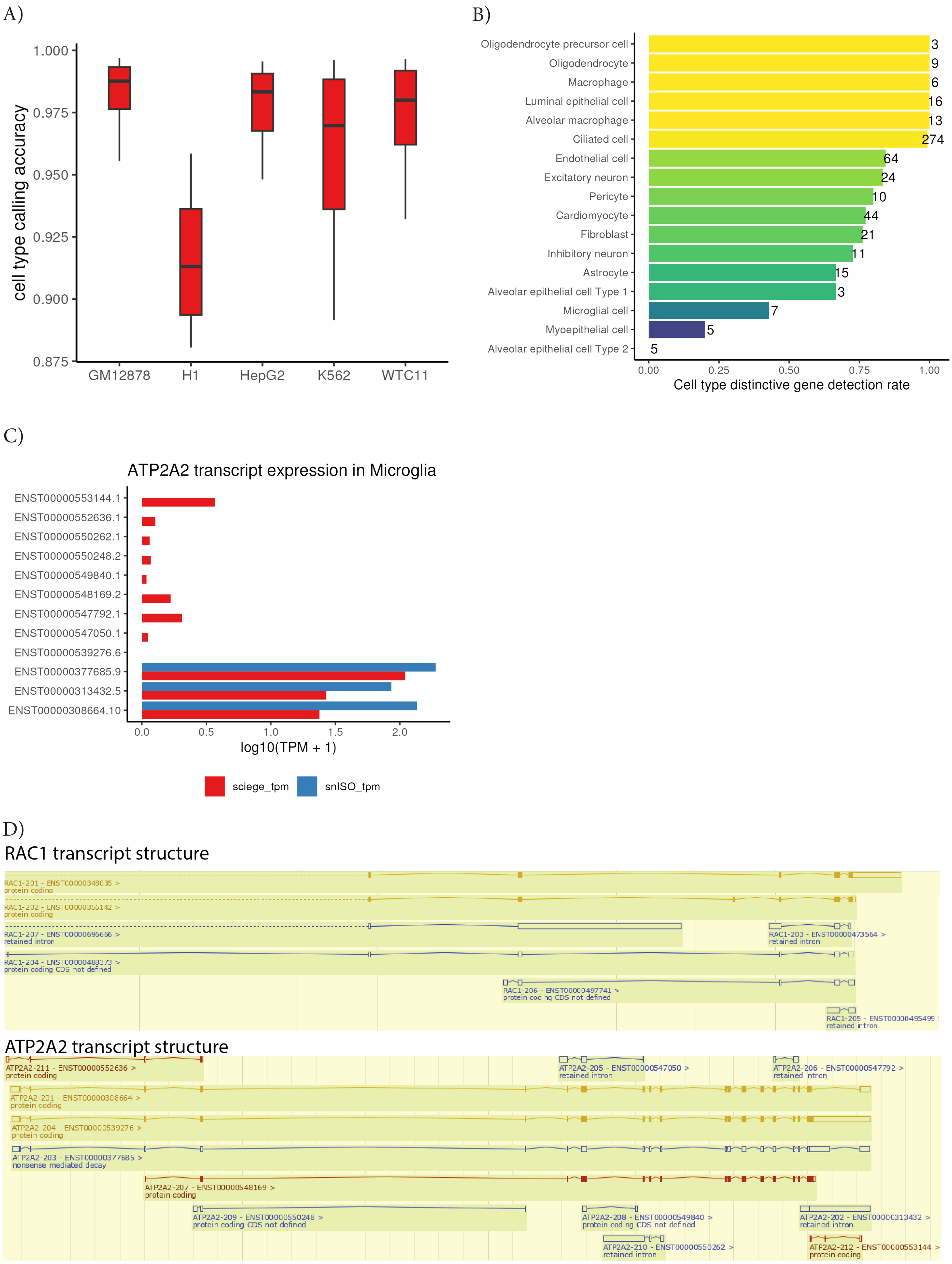


Figure S3: A) Boxplot showing accuracy of calling cell type specific isoforms in read-based simulation. X-axis represents names of cell lines. Y-axis represents accuracy of called cell type specific isoform in ground truth.Centerline of the box plot indicates median, box limit indicates first and third quartiles and points indicate outliers. B) Bar plot showing cell marker detection rate with Cell Marker 2.0. Colors and X-axis indicate detection rate. Numbers at the end of each bar indicate count of genes being tested. Y-axis indicates cell types. C) Barplot showing ATP2A transcript expression in Microglia. X-axis indicates log10(TPM + 1) value from either method. Blue color indicates TPM value from snISOrseq data while red color indicates Sciege estimate. Y-axis indicates transcript name. D) Edited screen shot of ensemble transcriot track for RAC1 and ATP2A2

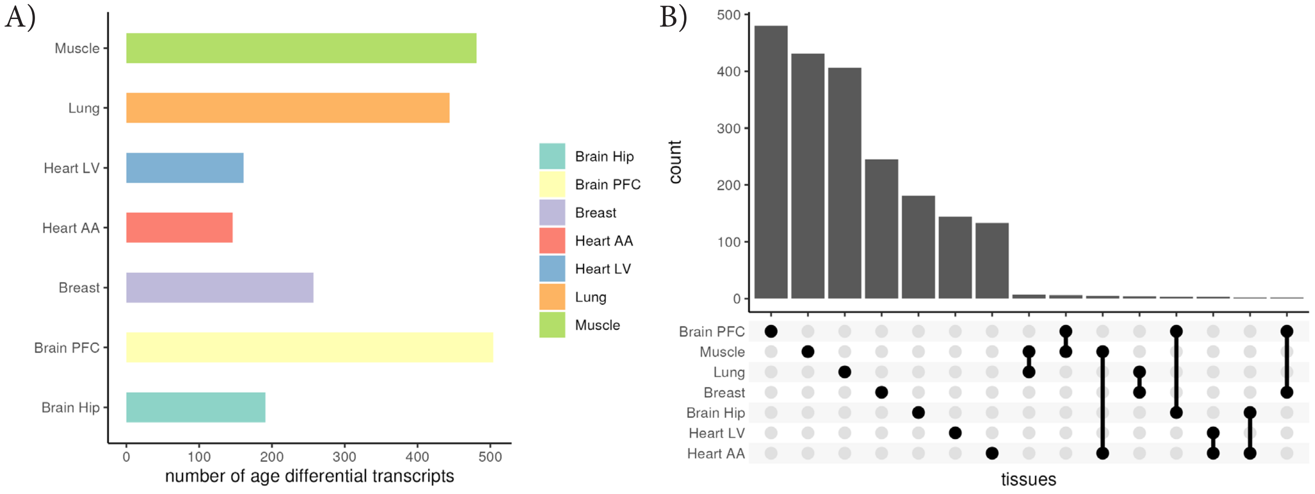


Figure S4: A) Barplot showing number of age differential transcripts per tissue. X-axis represents number of age differitial transcripts. Y-axis and color represents name of the tissue. B) Upset plot showing sharing of age differential transcrip between tissues. Top; barplot showing the number of age differential transcripts for each set. Bottom; Point plot showing which tissue repesents a set. Top of y-axis represents the number of age differential transcripts in each set. Bottom of y-axis represents the name of the tissue. X-axis represents set of sharing between tissues.

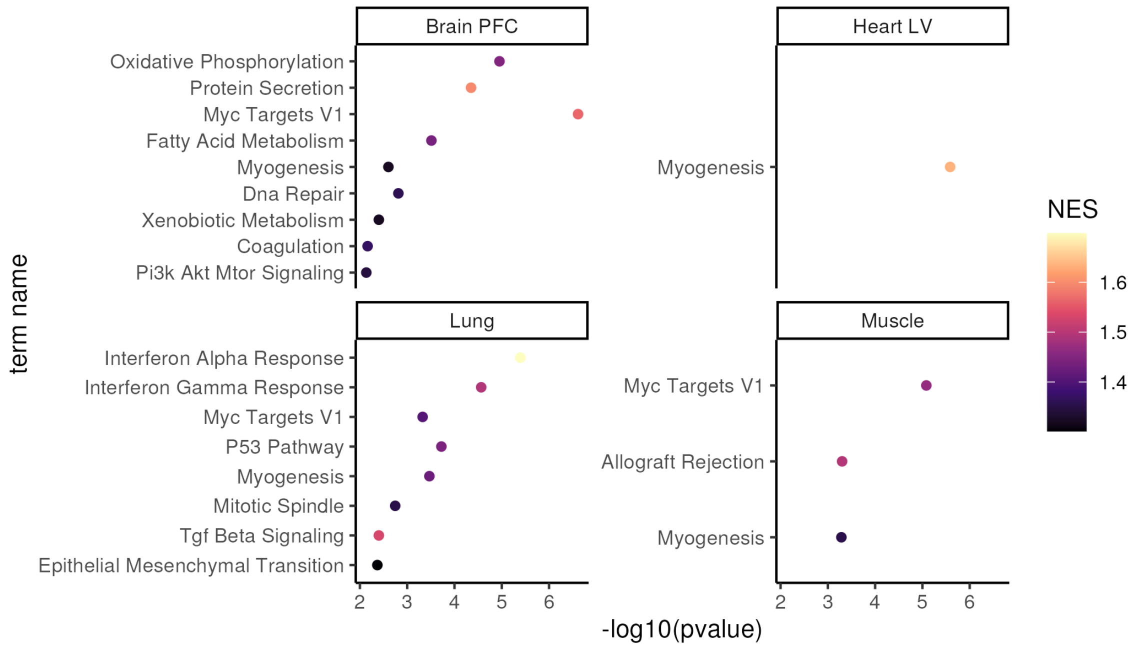


Figure S5: A) Dot plot showing Hallmark gene set enrichment of age differential transcript in four tissues. X-axis represents -log10 of p-value testing for enrichment for each gene set. Y-axis repesents the name of gene set. Color represents normalized enrichment score. Facet represents tissue names. Only FDR significant gene sets are shown.

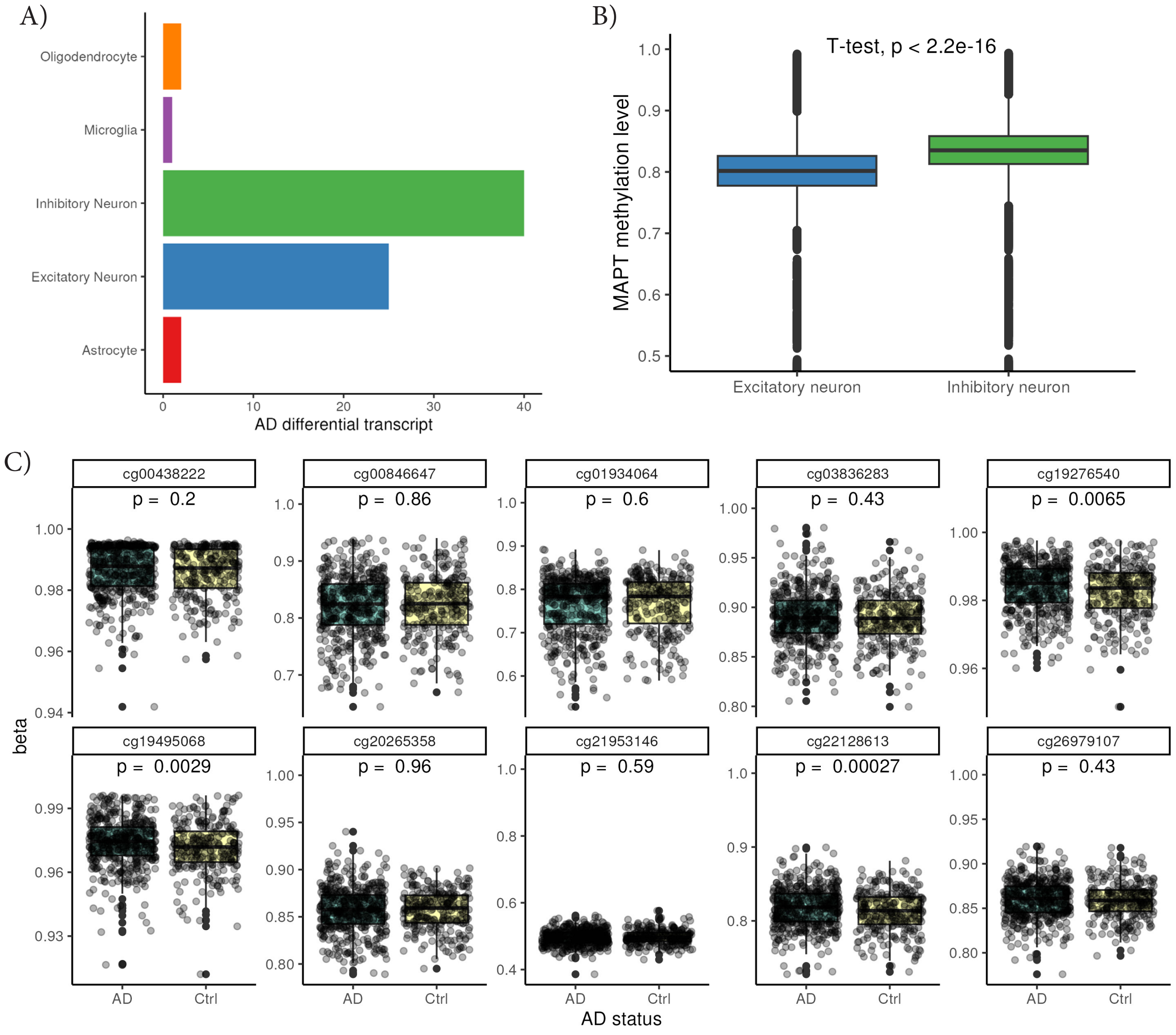


Figure S6: A) Barplot showing number of AD differential transcripts per cell type. X-axis represents number of AD differential transcripts. Y-axis and color represent cell type names. B) Box plot showing MAPT methylation distribution in two neuronal cell types. X-axis represents names of cell types. Y-axis represents MAPT methylation level. P-value from t-test is shown in top middle. Plot is cropeed from 0.5 to 1 on y-axis for visibiliry. Centerline of the box plot indicates median, box limit indicates first and third quartiles and points indicate outliers. C) Box plot showing methylation levels of CpG sites in CpG21. X-axis and color of box represent AD status. Y-axis represents methylation beta values. Facet represents the id of CpG site. P-values from linear model is shown in top middle (See Methods). Centerline of the box plot indicates median, box limit indicates first and third quartiles and points indicate outliers.

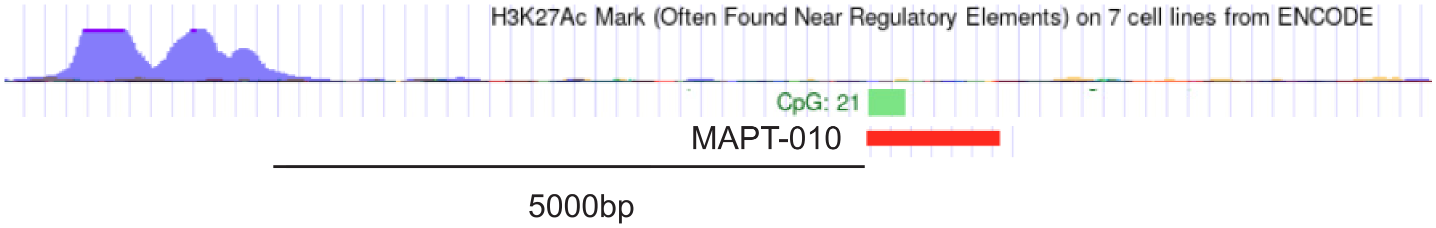


Figure S7: Modified screenshots of UCSC Genome Browser around MAPT-010 transcript. First track shows H327Ac mark from ENCODE. Second track shows CpG 21 island on MAPT transcript. Third track shows the body of MAPT-010 transcript.
